# Ghostbusting the national bird checklists: integrative evidence shows that *Pionus fuscus* does not occur in Colombia

**DOI:** 10.64898/2026.03.23.713821

**Authors:** Jhan C. Carrillo-Restrepo, Jorge Velásquez-Tibatá

## Abstract

Natural history collections underpin our understanding of species distributions, yet some historical records remain embedded in modern avifaunal checklists despite limited documentation and no independent verification. One such case concerns the Dusky Parrot *Pionus fuscus* in Colombia: although reported from specimens collected by Melbourne A. Carriker Jr. in 1942 in the Serranía de Perijá, the species has not been observed in the country for nearly eight decades yet continues to be included in national checklists and conservation assessments. We reassessed the validity of this record by applying a multi-evidence framework integrating historic archival reconstruction, specimen-based morphological comparisons, climatic niche analyses, biogeographic limit assessment and contemporary survey-effort data. Historical documentation and morphological evidence based on high-resolution specimen images and associated curatorial records demonstrate that the Carriker specimens correspond to *Pionus chalcopterus*, not *P. fuscus*. Climatic niche analyses reveal minimal environmental overlap between *P. chalcopterus* and *P. fuscus*, and place the Perijá locality within the climatic niche of *P. chalcopterus*, while regional biogeography and extensive modern birdwatching coverage provide no support for the occurrence of *P. fuscus* in Perijá. Together, these concordant lines of evidence demonstrate that *P. fuscus* does not occur in Colombia. Our findings support its removal from national bird lists and conservation assessments and highlight how integrated, multi-evidence reassessments of historical records strengthen ornithological baselines, improve biogeographic inference and ensure that conservation priorities rest on verifiable evidence.

## INTRODUCTION

Natural history collections represent an unparalleled record of Earth’s biodiversity. By preserving specimen-based evidence across space and time, they provide the empirical foundation for taxonomic revision, ecological and evolutionary research, and the delimitation of species ranges, thereby informing biogeographic inference and conservation planning (Ballard *et al*. 2017, Meineke *et al*. 2019, Nanglu *et al*. 2023). Despite the mobilization of museum databases to open data platforms and the rapid expansion of citizen science efforts (Sullivan 2009, Nelson & Ellis 2018), avian distributions, particularly in the Neotropics, remain incompletely resolved (Lees *et al*. 2020, Hughes *et al*. 2021). The continued documentation of new country records and unexpected range extensions underscores persistent gaps in our understanding of species range limits in one of the world’s most diverse regions (*e*.*g*., Lopes *et al*. 2009, Donegan 2012, Orihuela-Torres *et al*. 2020). These patterns highlight the need for sustained fieldwork and reaffirm the central role of museum specimens in documenting biodiversity, delineating species distributions and refining biogeographic and evolutionary interpretations (Meineke *et al*. 2019; Miller *et al*. 2020; Nanglu *et al*. 2023).

Historical vouchers and contemporary observations together underpin modern avian checklists and conservation assessments (Ballard *et al*. 2017). However, some early records were established under limited taxonomic frameworks and sparse documentation yet continue to be incorporated into current databases without independent verification (Marcer *et al*. 2022). When such records imply major distributional discontinuities, their persistence can propagate inaccuracies in ecological, evolutionary, and conservation analyses (Lees *et al*. 2020, Marcer *et al*. 2022). A case that illustrates this challenge concerns the reported occurrence of the Dusky Parrot (*Pionus fuscus*) in Colombia.

The Dusky Parrot is a psittacid species restricted to northeastern South America (Rodríguez-Mahecha & Hernández-Camacho 2002). Its range includes eastern Venezuela, the Guianas, and northern Brazil within the northern Amazon Basin, where it inhabits primarily lowland terra firme evergreen forests and, less frequently, seasonally flooded várzea and igapó, typically below 600 m (Collar *et al*. 2020a). A small, disjunct population has been suggested in the western slope of the Serranía de Perijá along the Venezuela–Colombia border, where it has been reported from montane forest (Ayerbe-Quiñones 2022, McMullan 2023).

The presence of *P. fuscus* in Colombia was first reported by Dugand (1948), who cited personal communication from Melbourne A. Carriker Jr., a major contributor to the documentation of Colombia’s avifauna through his extensive collecting efforts. According to Dugand, Carriker Jr. collected five individuals of *P. fuscus* at “Magdalena: Airoca, 1200 m, above Casacará, Sierra de Perijá (5 Carr.)”. Dugand further suggested that the toponym “Airoca” likely referred to “Hiroca, a Motilon village in the Sierra of Perijá mountains”. This report was notable because, when evaluated considering the species’ currently known distribution, it would represent an extension of approximately 950 km west of the nearest confirmed population in eastern Venezuela. Equally remarkable is the long-standing inclusion of this record in the literature. Since Dugand’s publication, *P. fuscus* has been included in nearly all major Colombian checklists (Meyer de Schauensee 1948-1952, Salaman *et al*. 2001, Donegan *et al*. 2016, Avendaño *et al*. 2017, Echeverry-Galvis *et al*. 2022) and field guides (Meyer de Schauensee 1966, Rodríguez-Mahecha & Hernández-Camacho 2002, Ayerbe-Quiñones 2022, McMullan 2023), thereby reinforcing the assumption that the species forms part of Colombia’s avifauna. It has even been included in the National Red List assessment as an Endangered species (Renjifo *et al*. 2014). The only notable exceptions are Hilty & Brown (1986, 2001) and Hilty (2021), who considered the record erroneous or possibly referring to an extirpated population.

Despite its long-standing acceptance, *P. fuscus* has not been observed or collected in Colombia since Dugand’s original account. Although survey coverage has been uneven across the broader Perijá region, particularly in the lowlands, ornithological work conducted over subsequent decades has not produced any independently verifiable record of the species (Donegan *et al*. 2003, López *et al*. 2014, Avendaño & Donegan 2015). The absence of modern records, combined with the ecological mismatch between the species’ known lowland habitats and the montane forests of Perijá, raises serious doubt about the validity of the historical report. Advances in ecological, morphological, and biogeographical analyses now allow historical records from natural history collections to be reassessed through the integration of multiple lines of evidence (Schindel & Cook 2018, Watanabe 2019, Miller *et al*. 2020). We therefore re-examined all available historical, morphological, and spatial data to evaluate whether the Colombian record of *P. fuscus* is accurate or instead represents a persistent error in the country’s ornithological literature. Beyond resolving the status of this species, our aim is to illustrate the value of multi-evidence reassessments for maintaining robust ornithological baselines.

## METHODS

### Historical documentation assessment

The documentary history associated with museum specimens often requires reconstruction from fragmentary archival sources. To clarify the basis of the Colombian record of *Pionus fuscus*, we revisited original publications, correspondence, field catalogues, and museum records linked to Carriker’s 1942 expedition to the Serranía de Perijá (Dugand 1948; Meyer de Schauensee 1948–1952; Paynter 1997). We examined Carriker’s handwritten field notes to identify specimen numbers, collection dates, and localities corresponding to the Perijá material. We also queried the National Museum of Natural History (NMNH) database to locate all *Pionus* specimens collected by Carriker Jr. in 1942 from the locality historically referred to as “Airoca/Eroca”.

### Morphological assessment

We assessed the taxonomic identity of the *Pionus* specimens collected by Carriker Jr. in 1942 in the Serranía de Perijá through detailed examination of high-resolution specimen images and associated curatorial records from the National Museum of Natural History (NMNH). Identifications were validated in consultation with NMNH curatorial staff and cross-checked against published diagnostic treatments of *Pionus fuscus* and *Pionus chalcopterus* (*e*.*g*., Collar *et al*. 2020a, 2020b). Assessment focused on species-level diagnostic characters, including head and neck coloration, orbital ring pigmentation, bill coloration, facial patterning, distribution of plumage coloration across upperparts and underparts, and overall chromatic contrast.

### Niche and biogeographic assessment

To evaluate whether independent climatic and biogeographic evidence supports the historical report of *Pionus fuscus* in Colombia, we compiled curated occurrence data for *P. fuscus* and *P. chalcopterus*, the morphologically most similar congener identified in our assessment. Records were downloaded from eBird (Sullivan *et al*. 2009) and cleaned following the protocol of Carrillo-Restrepo *et al*. (2025), which combines automated outlier detection with manual validation based on elevation, geographic distance, and spatial clustering.

Climatic niches for both species were characterized using 19 CHELSA bioclimatic variables at 30 arc-second (ca. 1 km) resolution (Karger *et al*. 2017). To reduce dimensionality and collinearity, we performed a joint principal component analysis (PCA) in environmental space using the function *espace_pca* in the *Wallace* package (Kass *et al*. 2023), based on environmental values extracted for cleaned occurrences and background points within each species’ accessible area (M; Barve *et al*. 2011). Accessible areas for each species were defined by intersecting validated occurrences with ecoregional boundaries (Dinerstein *et al*. 2017), and environmental predictors were restricted accordingly.

Niche overlap was quantified using two complementary approaches. First, a non-parametric kernel density framework implemented in *ecospat*.*niche*.*overlap* from the *ecospat* package (Di Cola *et al*. 2017) was used to calculate Schoener’s D (difference in probability densities between species) and Hellinger-based I (similarity based on niche overlap in environmental space) indices. Second, we constructed ellipsoidal-envelope ecological niche models (Farber & Kadmon 2003) for each species using the R package ntbox (Osorio-Olvera *et al*. 2020), following the methodological framework of Núñez-Penichet *et al*. (2021). Ellipsoids were defined by the centroid and covariance matrix of occupied environmental conditions, with limits determined from a chi-squared distribution of Mahalanobis distances (Etherington 2019), excluding 5% of marginal conditions. To incorporate variability, we generated 10 replicate ellipsoids per species using random subsamples comprising 75% of occurrence data, and final models were obtained by averaging centroids and covariance matrices across replicates. Overlap between species ellipsoids was quantified using the Jaccard index, calculated as the proportion of shared environmental space relative to the union of both ellipsoids. Statistical significance was assessed with a background-based null model (as in Núñez-Penichet *et al*. 2021), generating 1,000 pairs of ellipsoids from randomly sampled background points (sample size equal to empirical occurrences) and comparing observed overlap against the lower 5% of the null distribution. The georeferenced locality historically referred to as “Airoca/Eroca” was projected into the shared environmental space to evaluate its position relative to the estimated niches of both species.

Biogeographic plausibility was assessed by examining the spatial configuration of ecoregions and major habitat types between the core distribution of *P. fuscus* and the Serranía de Perijá using global (Dinerstein *et al*. 2017) and regional (Hazzi *et al*. 2018) frameworks, focusing on potential habitat discontinuities and physiographic barriers.

### Survey effort assessment

To determine whether the absence of modern records could reflect undersampling rather than true absence, we quantified contemporary birdwatching effort across northeastern Colombia and northwestern Venezuela using spatial summaries of eBird hotspots and complete checklists (Sullivan *et al*. 2009). Records of *P. fuscus* and *P. chalcopterus* were evaluated relative to survey intensity in the Perijá region.

## RESULTS

### Historical documentation evidence

The starting point for clarifying the Colombian record of *Pionus fuscus* was Dugand (1948), who reported several additions to the Colombian avifauna based on information communicated by M. A. Carriker Jr. through correspondence. In listing these records, Dugand noted that the supporting specimens were deposited “mainly in the Smithsonian Institution, Washington (USNM), and the Carnegie Museum, Pittsburgh (CM), with some in our Institute”, referring to the Instituto de Ciencias Naturales, Bogotá (ICN). However, he did not specify in which collection the specimens attributed to *P. fuscus* were deposited (Dugand 1948). This absence of primary specimen identifiers precluded direct verification of the material. Dugand (1948) also indicated that the locality name “Airoca” was likely a misspelling of “Hiroca”, highlighting that even the locality information required reinterpretation.

Decades later, Paynter (1997) re-examined Carriker’s field catalogues and addressed the toponymic inconsistencies. He pointed out a misprint “Eroca” in de Schauensee 1952a:1130 (Meyer de Schauensee 1948-1952) and noted that the presence of a “Quebrada Eroca” on IGAC maps of Cesar, approximately 25 km south of Agustín Codazzi, and the absence of “Hiroca” or “Airoca” on available cartography, suggested that “Eroca” was likely the correct spelling. Based on this clarification, we identified the most plausible location of the site at approximately 9°42’ N, 72°05’ W, at an elevation of ca. 1085 m, consistent with the altitudinal range reported by Carriker Jr. Paynter (1997) also documented the period of Carriker’s collections in the region, noting that he worked at “3,500–7,000 ft [1,050–2,125 m], 21–28, 30 Mar., 1–4, 6–10 Apr., and below Hiroca, at 2,000 ft [600 m], 11 May 1942 (USNM; CM, as ‘Eroca’)”. These clarifications established the spatial and temporal framework necessary to locate Carriker’s material and confirmed deposition in the United States National Museum (USNM), now referred to as the National Museum of Natural History (NMNH).

Building on Paynter’s work, we examined Carriker’s handwritten field notes from his 1942 Perijá expedition (Smithsonian Institution Archives, Record Unit 7297, Melbourne Armstrong Carriker Papers, Series 2, Folder 37). On page 37, line 8, we located the entry *“2326 Pionus fuscus Airoca, Apr. 6”*. Using this field number, we queried the NMNH specimen and associated digital records (Orrell & Informatics and Data Science Center - Digital Stewardship 2026) to locate all *Pionus* specimens collected by Carriker Jr. in 1942 from “Eroca”. This search returned six specimens collected between 26 March and 6 April 1942, catalogued as USNM 372620–372625 (Fig. 1). Notably, USNM-372624 carries field number 2326, establishing an unambiguous link between Carriker’s field number and the museum catalogue number (digital record available at http://n2t.net/ark:/65665/30a9193f1-f4e3-4c64-9705-6f2869915047, Orrell & Informatics and Data Science Center - Digital Stewardship 2026). This evidence also corrects a long-standing detail: although Dugand (1948) mentioned five individuals, NMNH records document six specimens (four males and two females), all identified as *Pionus chalcopterus* (Fig. 1), with collection dates and elevations consistent with those reported in the field documentation.

**Figure 1.**
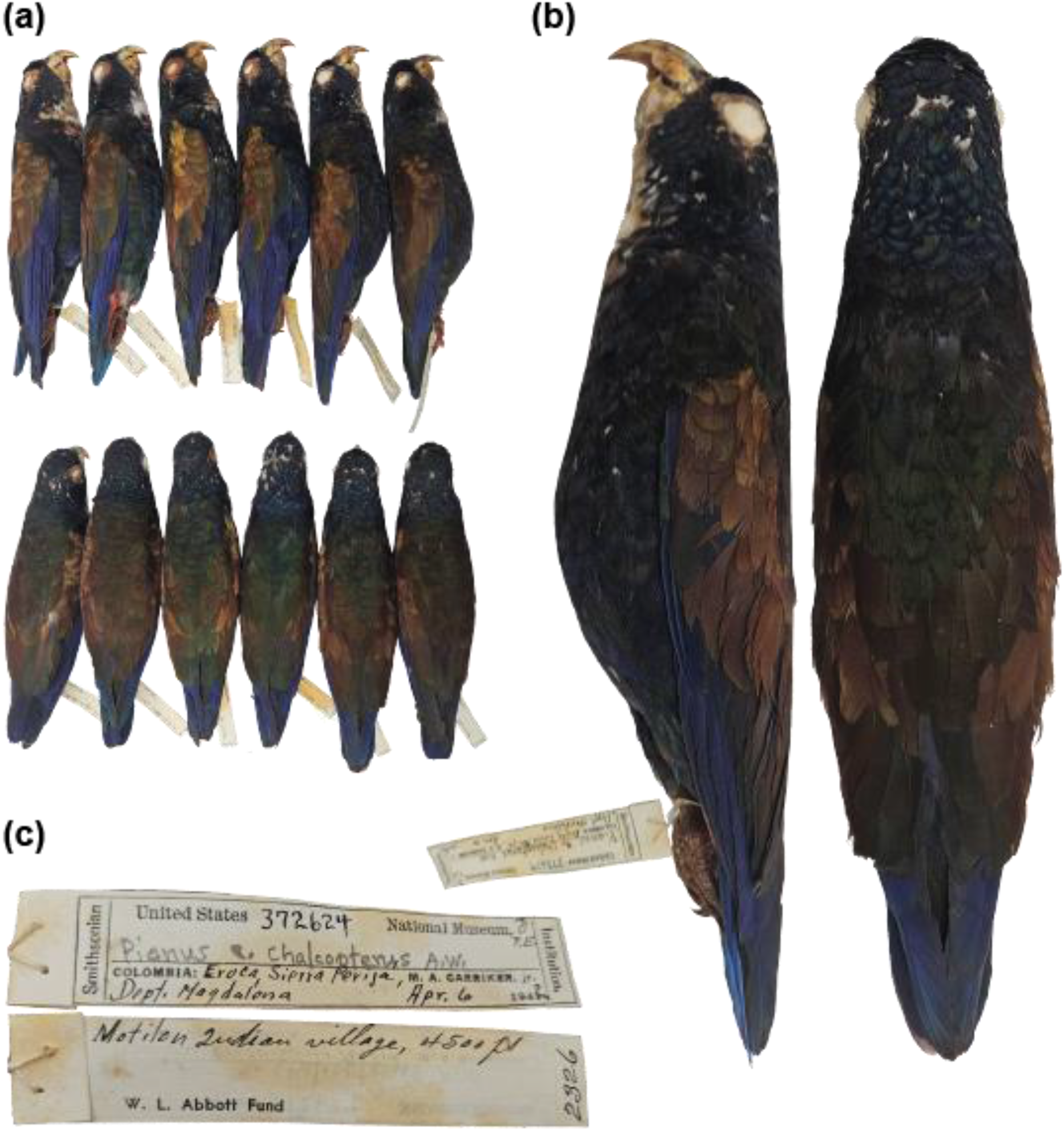
High-resolution images of specimens collected by Melbourne A. Carriker Jr. in the Serranía de Perijá in 1942. (a) Lateral and dorsal views of the six specimens deposited at the National Museum of Natural History (USNM 372620–372625), shown from left to right as: USNM-372620, USNM-372621, USNM-372622, USNM-372623, USNM-372624 and USNM-372625. All individuals are currently catalogued as *Pionus chalcopterus*. (b) Detailed view of specimen USNM-372624, which carries Carriker’s original field number 2326 and provides the direct link to the field note entry “2326 *Pionus fuscus* Airoca, Apr. 6”. Plumage characters visible in the specimen—including the bronze-sheened upperparts, dark bluish head and neck, flesh-coloured orbital ring and yellow bill—are consistent with *P. chalcopterus* and inconsistent with *P. fuscus*. (c) Original labels of specimen USNM-372624 indicating a male collected at “Eroca, Sierra Perijá, Dept. Magdalena” on 6 April 1942 by M. A. Carriker Jr.; additional information on the reverse indicates “Motilon Indian Village, 4500 ft” (≈1500 m). These archival data confirm the locality and date associated with the specimen series examined in this study. Photographs by C. Milensky, Smithsonian Institution, National Museum of Natural History.

### Morphological evidence

Examination of high-resolution images of the museum skins, together with published diagnostic treatments and comparative illustrations (Collar *et al*. 2020a, 2020b), revealed plumage characters consistent with *Pionus chalcopterus* (Fig. 1-2). The specimens exhibit a bronze-brown to very dark navy-blue head and neck, a flesh-colored orbital ring, and a yellow bill. The chin is flecked with white above a pink, scaly bib that transitions into deep blue underparts. The back, mantle, scapulars, and wing coverts display a distinct bronze sheen, whereas the rump, flight feathers, and tail are uniformly deep blue (Fig. 1-2; Collar *et al*. 2020b). For comparison, published descriptions of material of *P. fuscus* indicate a slaty-blue head, a pale bluish-grey orbital ring, and a small red patch below the nares. The face is bordered by a subtle whitish fringe; the upperparts are dark brown with paler feather margins; the underparts range from dark greyish to reddish chocolate, often with diffuse barring; and the wings and tail are dark blue, giving the species a darker overall appearance (Fig. 1-2; Collar *et al*. 2020a). None of these diagnostic characters observed in *P. fuscus* were present in the specimens collected by Carriker Jr. at “Eroca” in the Serranía de Perijá. Instead, all examined individuals exhibit the suite of plumage traits described for *P. chalcopterus*. Consistent with these observations, NMNH curatorial records catalogue all specimens under *Pionus chalcopterus*, despite their original annotation as *P. fuscus* in Carriker’s field notes. The moment or circumstances under which this taxonomic correction occurred are not documented.

**Figure 2.**
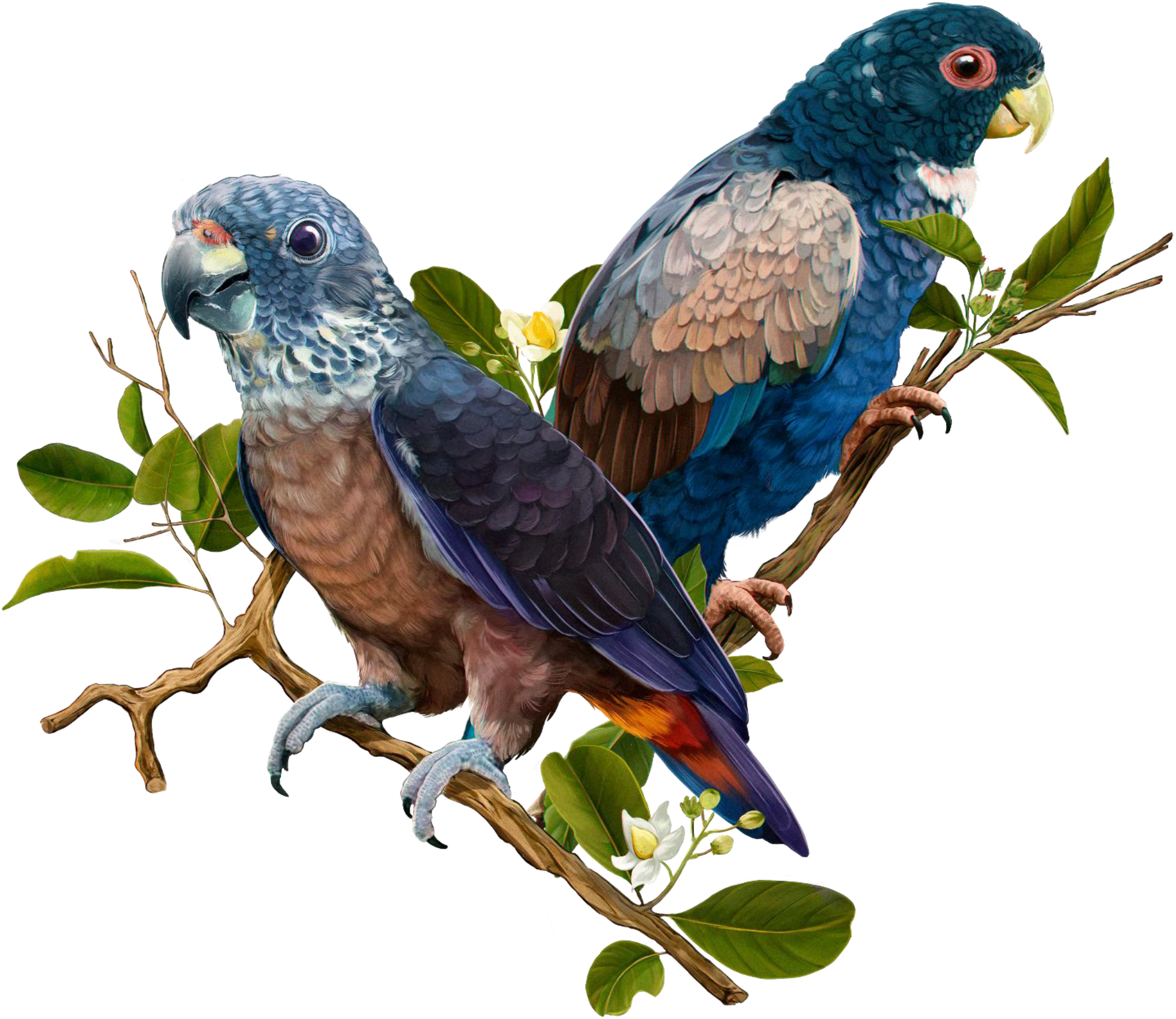
Comparative illustration highlighting morphological and habitat differences between the focal species. This illustration contrasts *Pionus fuscus* (left) and *Pionus chalcopterus* (right) and depicts each perched on representative host plants. *P. fuscus* appears on *Eschweilera subglandulosa*, typical of lowland forests, while *P. chalcopterus* is shown on the Colombian montane oak (*Trigonobalanus excelsa*), a species endemic to montane forests. Illustration by Steven Pinzon.

### Climatic niche and biogeographical evidence

After cleaning and validation, a total of 725 occurrence records of *Pionus fuscus* and 4,396 of *Pionus chalcopterus* were retained for analysis (from initial datasets of 6,969 and 88,404 records, respectively). These curated occurrences were used to characterize climatic niches and quantify niche overlap between both taxa. The joint PCA analysis of 19 CHELSA bioclimatic variables summarized the environmental variation occupied by both species, with the first three axes explaining 85.52% of total variance. Within this reduced environmental space, niche overlap estimated using the kernel-density framework (ecospat, Di Cola *et al*. 2017) was extremely low (Schoener’s D = 0.012; Hellinger-based I = 0.071), indicating minimal similarity in occupied climatic conditions. Ellipsoidal-envelope niche models yielded consistent results. The observed Jaccard overlap between ellipsoids of *P. fuscus* and *P. chalcopterus* was J = 0.24, a value below the lower 5% confidence limit of the null distribution generated from background-based simulations (null threshold = 0.287). Projection of the georeferenced locality historically referred to as “Airoca/Eroca” into this environmental space placed it within the ellipsoid of *P. chalcopterus* and outside that of *P. fuscus* (Fig. 3).

**Figure 3.**
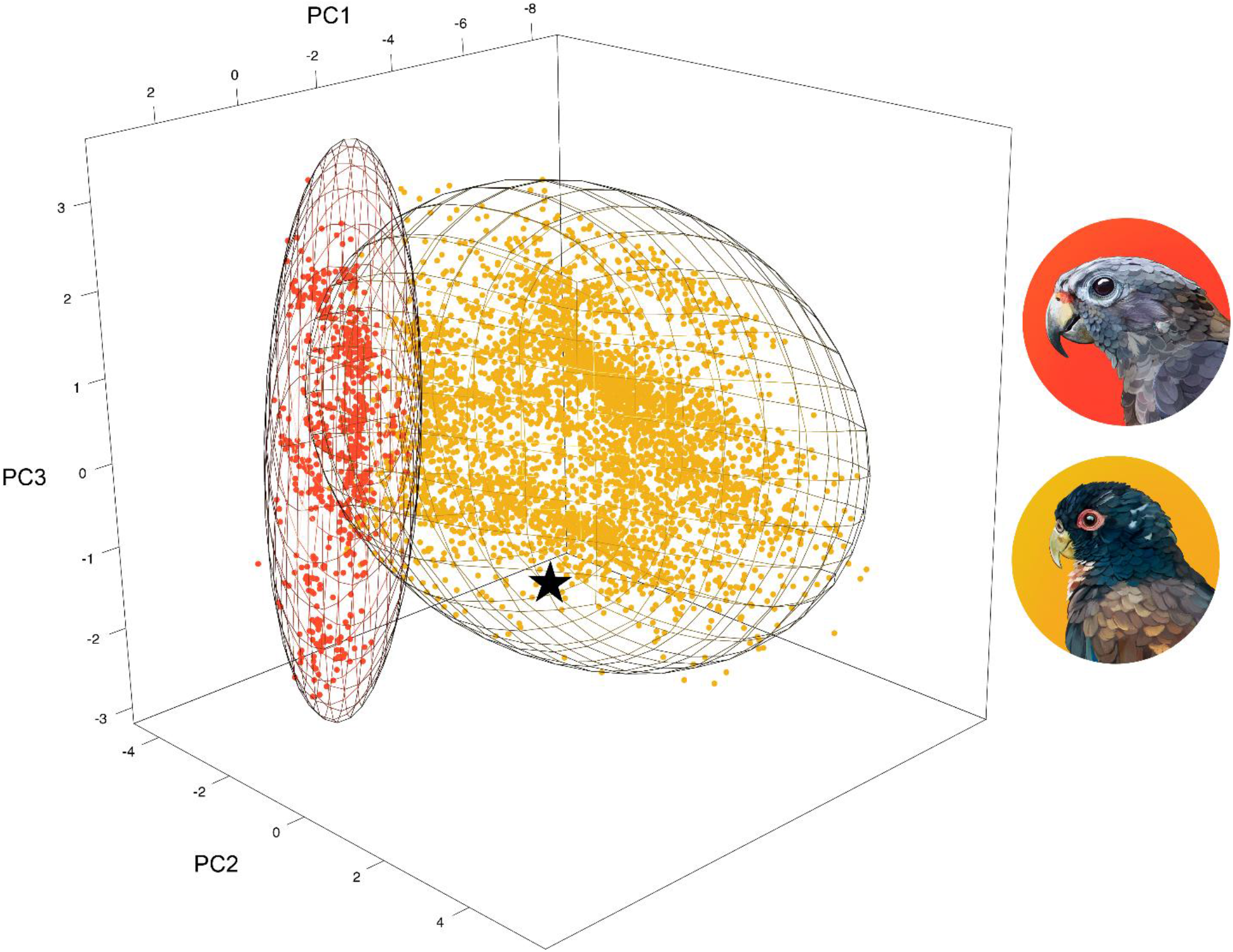
Climatic niche differentiation between the focal species. Three-dimensional climatic niche space defined by the first three principal components (PC1–PC3) derived from 19 CHELSA bioclimatic variables. The red ellipsoid represents the modeled climatic niche of *Pionus fuscus*, and the yellow ellipsoid that of *Pionus chalcopterus*, estimated using parametric 3D ellipsoids (ntbox). The black star marks Carriker’s 1942 “Eroca” specimens, all of which fall inside the climatic niche of *P. chalcopterus* and outside that of *P. fuscus*. Illustrations by Steven Pinzon.

Examination of ecoregional frameworks revealed a marked spatial discontinuity between the core distribution of *P. fuscus* in the Guianan Shield and the Serranía de Perijá (Fig. 4a). The intervening landscape is characterized by extensive dry forest formations, savanna systems, and high-elevation Andean environments (Dinerstein *et al*. 2017, Hazzi *et al*. 2018), contrasting with the humid lowland forest habitats where *P. fuscus* is documented.

**Figure 4.**
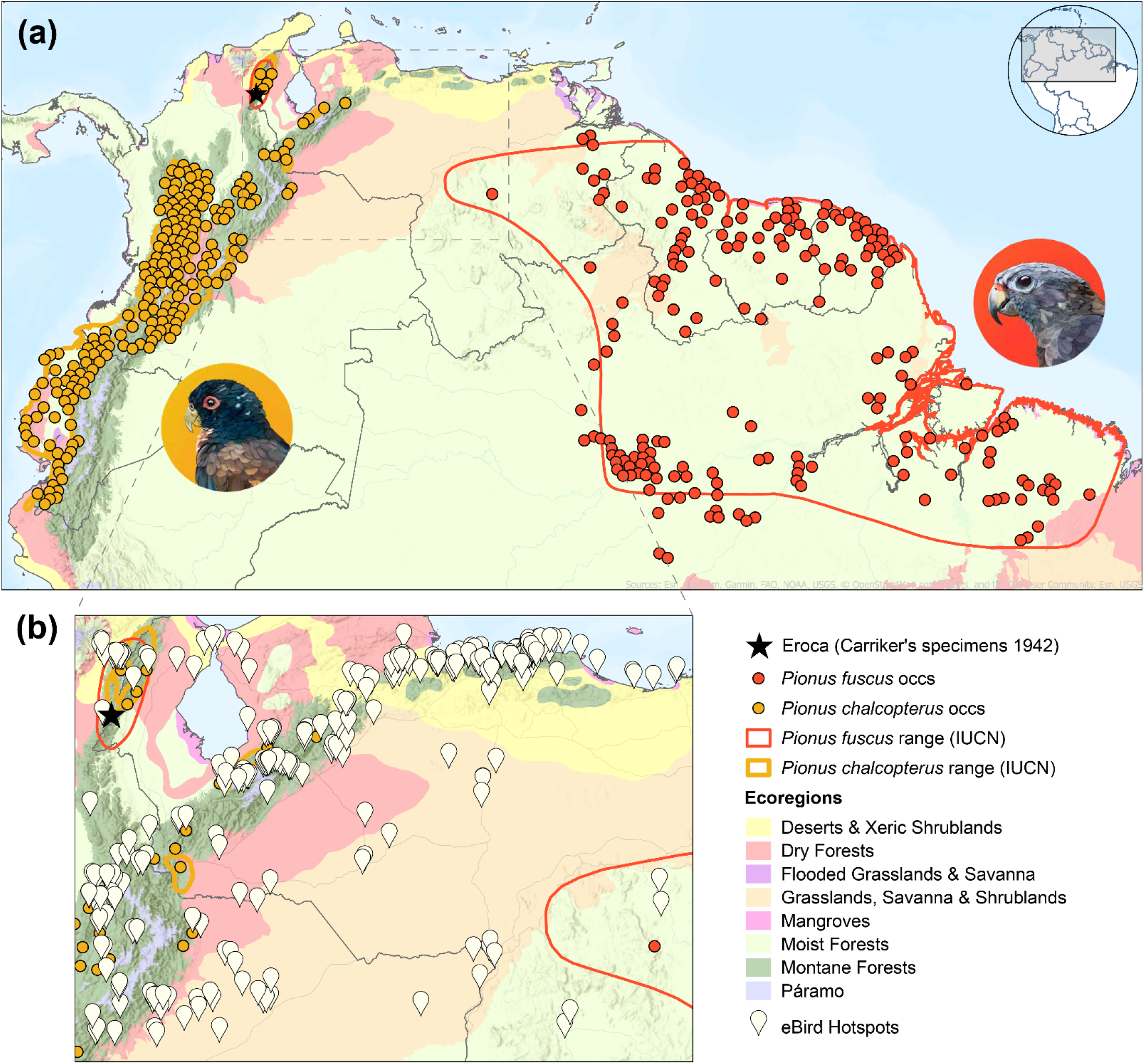
Biogeographical limits and sampling evidence regarding the occurrence of the focal species in the Serranía de Perijá. (a) Biogeographical comparison of the distributional limits of *Pionus fuscus* (red) and *Pionus chalcopterus* (yellow), highlighting that only the latter occupies the montane forests of Perijá. (b) Summary of birdwatching effort across northwestern Venezuela and northeastern Colombia, showing extensive sampling coverage and consistent documentation of *P. chalcopterus* in Perijá, with *P. fuscus* remaining entirely undetected. Terrestrial ecoregions based on Dinerstein *et al*. (2017).

### Survey-effort evidence

Finally, contemporary sampling data provides an additional independent line of evidence. Across northeastern Colombia and northwestern Venezuela, eBird currently includes 1,499 active hotspots, 50 of which contain more than 100 complete checklists and over 300 recorded species, reflecting sustained survey coverage in the region (Fig. 4b). Despite this sampling intensity, *Pionus fuscus* has not been reported from the Perijá region. In contrast, *P. chalcopterus* has been documented in 1,981 complete checklists, and all 58 records supported by photographic evidence correspond to this species.

## DISCUSSION

All lines of evidence examined in this study, historical, morphological, climatic, biogeographical, and survey effort, converge on a single conclusion: the Dusky Parrot (*Pionus fuscus*) does not occur in Colombia. The historical record reported by Dugand (1948) was based on a misidentified series of *P. chalcopterus* collected by Carriker Jr. in 1942, and no independent evidence supports the presence of *P. fuscus* in the country.

Archival reconstruction linked Carriker’s field notes to six specimens deposited at NMNH, and morphological examination showed that all exhibit the diagnostic plumage characters of *P. chalcopterus* (Collar *et al*. 2020a, 2020b). Although fading associated with specimen age can affect superficial coloration in museum skins, the visible structural and chromatic traits are consistent with *P. chalcopterus* and incompatible with *P. fuscus* (Fig. 1-2). The convergence of documentary and morphological evidence resolves a misidentification that persisted in Colombian literature for more than seven decades and removes the only historical basis for including *P. fuscus* in the Colombian avifauna.

Independent ecological analyses reinforce this conclusion. Kernel-based and ellipsoidal niche models revealed minimal climatic overlap between *P. fuscus* and *P. chalcopterus*, and the 1942 locality falls within the climatic niche of *P. chalcopterus* but outside that of *P. fuscus* (Fig. 3). Biogeographic structure provides an additional constraint (Fig. 4a). *Pionus fuscus* is restricted to lowland humid forests of the Guianan Shield and northern Brazil, typically below 600 m. Reaching the Serranía de Perijá would require crossing the Apure–Villavicencio and Maracaibo dry forests, the Llanos savannas, and the high-elevation systems of the Cordillera de Mérida—landscapes that differ markedly from its known habitat and form a large ecological discontinuity between the Guianan Shield and Perijá (Fig. 4a; Dinerstein *et al*. 2017, Hazzi *et al*. 2018). In contrast, the environmental conditions of Perijá fall within the documented range of *P. chalcopterus* (Fig. 3). Together, climatic differentiation and regional landscape configuration render the occurrence of *P. fuscus* in Perijá highly implausible.

Contemporary survey data provides a further independent test. Although the broader Perijá region has been surveyed repeatedly, sampling coverage of the lowlands has been more limited, and much of that landscape has long been heavily deforested (Donegan *et al*. 2003, López *et al*. 2014, Avendaño & Donegan 2015), leaving little suitable habitat for a lowland humid-forest species such as *P. fuscus*. Even so, *P. fuscus* has not been reported from the region, whereas *P. chalcopterus* is regularly documented (Fig. 4b). The rediscovery of other lowland forest birds in remnant habitats of northern Colombia, such as *Crypturellus saltuarius* (Donegan *et al*. 2003, Laverde-R. & Cadena 2014), shows that the absence of records alone is not always decisive. However, such cases typically involve cryptic or low-density taxa capable of persisting undetected in small relict populations. In contrast, *Pionus* parrots are conspicuous, vocal, and often occur in detectable groups, making prolonged non-detection under survey effort and expanding birdwatching coverage unlikely. The absence of modern records therefore aligns with all other lines of evidence.

This reassessment has immediate implications for national avifaunal lists. Under the criteria of the Comité Colombiano de Registros Ornitológicos (https://ccro.asociacioncolombianadeornitologia.org/criterios/), species should be included only when supported by independently verifiable evidence (*e*.*g*. catalogued museum specimens, photographic or audio vouchers with reliable metadata, GPS-tracked individuals, or peer-reviewed publications). In the case of *P. fuscus*, no such evidence exists for Colombia, and the Carriker Jr. series has been shown to belong entirely to *P. chalcopterus* (Fig. 1). We therefore recommend that *P. fuscus* be formally removed from Colombian bird lists.

This recommendation should be viewed within a broader recent trend in Colombian ornithology. Over the past two decades, the national bird list has grown substantially through new field discoveries, taxonomic revisions and improved coverage of previously underexplored regions (Donegan 2012, Avendaño *et al*. 2017, Echeverry-Galvis *et al*. 2022), but parallel efforts have also sought to strengthen the evidentiary standards by which species are accepted. Critical revisions of the Colombian avifauna have already resulted in the removal of several taxa based on doubtful documentation or misidentified material (e.g. Donegan *et al*. 2009, 2010, Lobo & Henriquez 2014, Echeverry-Galvis *et al*. 2022). Our reassessment of *P. fuscus* contributes to this ongoing process of refining the national list through more rigorous evaluation of legacy records.

The correction also carries ecological and conservation implications. *P. fuscus* was listed as Endangered in the *Libro Rojo de Aves de Colombia* (Renjifo *et al*. 2014) and incorporated into national threatened-species legislation (Resolution 1912 of 2017, Ministry of Environment and Sustainable Development, Republic of Colombia). Maintaining species under threat categories without confirmed national presence risks misdirecting research priorities, conservation planning, and funding away from taxa with verified occurrence and demonstrated need. Accurate species lists are foundational to effective biodiversity policy. More broadly, removing *P. fuscus* from the Colombian avifauna prevents its erroneous inclusion in biogeographical and evolutionary analyses using widely used distributional databases such as BirdLife International & Handbook of the Birds of the World (2026) range maps. Spurious locality data can bias inferences about range limits, historical connectivity, and diversification processes, particularly in groups such as Neotropical parrots where distributional boundaries inform phylogeographic hypotheses (*e*.*g*. Ribas *et al*. 2007).

This case underscores the importance of revisiting legacy records through integrated approaches that combine archival research, morphological validation, ecological modelling, and contemporary survey data. Natural history collections remain indispensable to biodiversity science, but their value depends on continued taxonomic scrutiny, curatorial precision, and ecological validation to maintain reliable ornithological and biogeographical baselines.

## ACKNOWLEDGMENTS

We acknowledge eBird and GBIF for their commitment to maintaining open-access biodiversity data, which made our biogeographical and survey-effort analyses possible; GBIF also provided access to digitized museum records that were essential for the historical component of this study. We thank A. M. Cuervo and the ornithological collection of the Instituto de Ciencias Naturales (ICN), Universidad Nacional de Colombia, for granting access to archival materials that allowed us to examine correspondence between M. A. Carriker Jr. and A. Dugand regarding the inclusion of *Pionus fuscus* in Colombia. We also thank R. G. Gilreath (Smithsonian Institution Archives) for generously scanning the folder containing Carriker’s handwritten field notes, which proved essential for resolving this longstanding mystery. We are also grateful to C. Milensky (Smithsonian Institution, National Museum of Natural History) for providing high quality photographs of the Carriker’s specimens housed in the NMNH collection. We are grateful to Steven Pinzón (BAHLA Estudio Creativo) for creating the outstanding *Pionus* illustrations specifically for this study. Finally, we thank T. M. Donegan for helpful comments and suggestions that improved the manuscript.

## DATA AVAILABILITY STATEMENT

The data and R code supporting the findings of this study are openly available in Zenodo at https://doi.org/10.5281/zenodo.19198462. The official Spanish translation of the manuscript is likewise deposited in this repository (Carrillo-Restrepo & Velásquez-Tibatá 2026).

## REFERENCES

Avendaño, J.E. & Donegan, T.M. 2015. A distinctive new subspecies of Scytalopus griseicollis (Aves, Passeriformes, Rhinocryptidae) from the northern Eastern Cordillera of Colombia and Venezuela. ZooKeys 506: 137–153. 10.3897/zookeys.506.9553

Avendaño, J.E., Bohórquez, C.I., Rosselli, L., Arzuza-Buelvas, D., Estela, F.A., Cuervo, A.M., Stiles, F.G. & Renjifo, L.M. 2017. Lista de chequeo de las aves de Colombia: una síntesis del estado del conocimiento desde Hilty & Brown (1986). Ornitol. Colomb. 16: 1–83.

Ayerbe-Quiñones, F. 2022. Guía ilustrada de la avifauna colombiana. Wildlife Conservation Society– Colombia, Bogotá, Colombia.

Ballard, H.L., Robinson, L.D., Young, A.N., Pauly, G.B., Higgins, L.M., Johnson, R.F. & Tweddle, J.C. 2017. Contributions to conservation outcomes by natural history museum-led citizen science: examining evidence and next steps. Biol. Conserv. 208: 87–97. 10.1016/j.biocon.2016.08.040

Barve, N., Barve, V., Jiménez-Valverde, A., Lira-Noriega, A., Maher, S.P., Peterson, A.T. & Villalobos, F. 2011. The crucial role of the accessible area in ecological niche modeling and species distribution modeling. Ecol. Model. 222: 1810–1819. 10.1016/j.ecolmodel.2011.02.011

BirdLife International & Handbook of the Birds of the World. 2026. Bird species distribution maps of the world. Version 2026.1. BirdLife International, Cambridge, UK. Available at http://datazone.birdlife.org/species/requestdis (accessed 2026).

Carrillo-Restrepo, J.C., Linero-Triana, D., Herzog, S.K. & Velásquez-Tibatá, J.I. 2025. Area of habitat maps and validated occurrences for neotropical birds of conservation concern. Sci. Data 12: 1108. 10.1038/s41597-025-05393-y

Carrillo-Restrepo, J. C. & Velásquez-Tibatá, J. 2026. Data from: Ghostbusting the national bird checklist: integrative evidence shows that Pionus fuscus does not occur in Colombia. Zenodo. 10.5281/zenodo.19198462

Collar, N.,Boesman, P.F.D. & Kirwan, G.M. 2020a. Dusky Parrot (Pionus fuscus), version 1.0. In del Hoyo, J., Elliott, A., Sargatal, J., Christie, D.A. & de Juana, E. (eds) Birds of the World. Cornell Lab of Ornithology, Ithaca, NY, USA. 10.2173/bow.duspar1.01

Collar, N., Boesman, P.F.D. & Kirwan, G.M. 2020b. Bronze-winged Parrot (Pionus chalcopterus), version 1.0. In del Hoyo, J., Elliott, A., Sargatal, J., Christie, D.A. & de Juana, E. (eds) Birds of the World. Cornell Lab of Ornithology, Ithaca, NY, USA. 10.2173/bow.brwpar1.01

Di Cola, V., Broennimann, O., Petitpierre, B., Breiner, F.T., D’Amen, M., Randin, C. & Guisan, A. 2017. ecospat: an R package to support spatial analyses and modelling of species niches and distributions. Ecography 40: 774–787. 10.1111/ecog.02671

Dinerstein, E., Olson, D., Joshi, A., Vynne, C., Burgess, N.D., Wikramanayake, E. & Saleem, M. 2017. An ecoregion-based approach to protecting half the terrestrial realm. BioScience 67: 534–545. 10.1093/biosci/bix014

Donegan, T.M., Huertas, B.C. & Briceño, E.R. 2003. Status of the Magdalena Tinamou Crypturellus saltuarius in the type locality and surrounding lower Magdalena Valley. Cotinga 19: 34–39.

Donegan, T.M., Salaman, P.G.W. & Caro, D. 2009. Revision of the status of various bird species occurring or reported in Colombia. Conserv. Colomb. 8: 80–86.

Donegan, T.M., Salaman, P.G.W., Caro, D. & McMullan, M. 2010. Revision of the status of bird species occurring in Colombia 2010. Conserv. Colomb. 13: 25–54.

Donegan, T.M. 2012. Range extensions and other notes on the birds and conservation of the Serranía de San Lucas, an isolated mountain range in northern Colombia. Bull. Br. Ornithol. Club 132: 140–161.

Donegan, T., Verhelst, J.C., Ellery, T., Cortés-Herrera, O. & Salaman, P.G.W. 2016. Revision of the status of bird species occurring or reported in Colombia 2016 and assessment of BirdLife International’s new parrot taxonomy. Conserv. Colomb. 24: 12–36.

Dugand, A. 1948. Notas ornitológicas colombianas, IV. Caldasia 5: 157–199.

Echeverry-Galvis, M.Á., Acevedo-Charry, O., Avendaño, J.E., Gómez, C., Stiles, F.G., Estela, F.A. & Cuervo, A.M. 2022. Lista oficial de las aves de Colombia 2022: adiciones, cambios taxonómicos y actualizaciones de estado. Ornitol. Colomb. 22: 25–51.

Etherington, T.R. 2019. Mahalanobis distances and ecological niche modelling: correcting a chi-squared probability error. PeerJ 7: e6678. 10.7717/peerj.6678

Farber, O. & Kadmon, R. 2003. Assessment of alternative approaches for bioclimatic modelling with special emphasis on the Mahalanobis distance. Ecol. Model. 160: 115–130. 10.1016/S0304-3800(02)00327-7

Hazzi, N.A., Moreno, J.S., Ortiz-Movliav, C. & Palacio, R.D. 2018. Biogeographic regions and events of isolation and diversification of the endemic biota of the tropical Andes. Proc. Natl Acad. Sci. USA 115: 7985–7990. 10.1073/pnas.1803908115

Hilty, S.L. & Brown, W.L. 1986. A guide to the birds of Colombia. Princeton University Press, Princeton, NJ.

Hilty, S.L. & Brown, W.L. 2001. Guía de las aves de Colombia. American Bird Conservancy, Universidad del Valle & Sociedad Antioqueña de Ornitología, Cali.

Hilty, S.L. 2021. Birds of Colombia. Lynx Edicions, Barcelona.

Hughes, A.C., Orr, M.C., Ma, K., Costello, M.J., Waller, J., Provoost, P., Yang, Q., Zhu, C. & Qiao, H. 2021. Sampling biases shape our view of the natural world. Ecography 44: 1259–1269. 10.1111/ecog.05926

Karger, D.N., Conrad, O., Böhner, J., Kawohl, T., Kreft, H., Soria-Auza, R.W., Zimmermann, N.E., Linder, H.P. & Kessler, M. 2017. Climatologies at high resolution for the earth’s land surface areas. Sci. Data 4: 170122. 10.1038/sdata.2017.122

Kass, J.M., Pinilla-Buitrago, G.E., Paz, A., Johnson, B.A., Grisales-Betancur, V., Meenan, S.I., Attali, D., Broennimann, O., Galante, P.J., Maitner, B.S., Owens, H., Varela, S., Aiello-Lammens, M., Merow, C., Blair, M.E. & Anderson, R.P. 2023. Wallace 2: a shiny app for modelling species niches and distributions redesigned to facilitate expansion via module contributions. Ecography e06547. 10.1111/ecog.06547

Laverde-R. O. & Cadena, C.D. 2014. Taxonomy and conservation: a tale of two tinamou species groups (Tinamidae, Crypturellus). J. Avian Biol. 45: 484–492. 10.1111/jav.00298

Lees, A.C., Rosenberg, K.V., Ruiz-Gutierrez, V., Marsden, S., Schulenberg, T.S. & Rodewald, A.D. 2020. A roadmap to identifying and filling shortfalls in Neotropical ornithology. Auk 137: ukaa048. 10.1093/auk/ukaa048

Lobo, Y. & Henriques, J.C. 2014. Cave Swallow Petrochelidon fulva and Couch’s Kingbird Tyrannus couchii: a discussion of two difficult cases of potential records for Colombia based on museum specimens. Conserv. Colomb. 21: 58–62.

Lopes, L.E., Malacco, G.B., Alteff, E.F., Vasconcelos, M.F., Hoffmann, D. & Silveira, L.F. 2010. Range extensions and conservation of some threatened or little known Brazilian grassland birds. Bird Conserv. Int. 20: 84–94. 10.1017/S0959270909990190

López, J.P., Avendaño, J.E., Gutiérrez-Pinto, N. & Cuervo, A.M. 2014. The birds of the Serranía de Perijá: the northernmost avifauna of the Andes. Ornitol. Colomb. 14: 62–93.

Marcer, A., Chapman, A.D., Wieczorek, J.R., Picó, F.X., Uribe, F., Waller, J. & Ariño, A.H. 2022. Uncertainty matters: ascertaining where specimens in natural history collections come from and its implications for predicting species distributions. Ecography e06025. 10.1111/ecog.06025

McMullan, M. 2023. Guía de campo de las aves de Colombia. McMullan Birding & Publishers, Cali.

Meineke, E.K., Davies, T.J., Daru, B.H. & Davis, C.C. 2019. Biological collections for understanding biodiversity in the Anthropocene. Phil. Trans. R. Soc. B 374: 20170386. 10.1098/rstb.2017.0386

Meyer de Schauensee, R. 1948–1952. The birds of the Republic of Colombia. Caldasia 22–26: 251–1212.

Meyer de Schauensee, R. 1966. The species of birds of South America with their distribution. Livingston Press, Narbeth, PA.

Miller, S.E., Barrow, L.N., Ehlman, S.M., Goodheart, J.A., Greiman, S.E., Lutz, H.L. & Light, J.E. 2020. Building natural history collections for the twenty-first century and beyond. BioScience 70: 674–687. 10.1093/biosci/biaa069

Nanglu, K., de Carle, D., Cullen, T.M., Anderson, E.B., Arif, S., Castañeda, R.A. & Astudillo-Clavijo, V. 2023. The nature of science: the fundamental role of natural history in ecology, evolution, conservation and education. Ecol. Evol. 13: e10621. 10.1002/ece3.10621

Nelson, G. & Ellis, S. 2019. The history and impact of digitization and digital data mobilization on biodiversity research. Phil. Trans. R. Soc. B 374: 20170391. 10.1098/rstb.2017.0391

Nuñez-Penichet, C., Cobos, M.E. & Soberón, J. 2021. Non-overlapping climatic niches and biogeographic barriers explain disjunct distributions of continental Urania moths. Front. Biogeogr. 13. 10.21425/F5FBG52142

Orrell, T. & Informatics and Data Science Center - Digital Stewardship. 2026. NMNH extant specimen records (USNM, US). National Museum of Natural History, Smithsonian Institution. Occurrence dataset. 10.15468/hnhrg3 accessed via GBIF.org on [12 march 2026].

Orihuela-Torres, A., Tinoco, B., Ordóñez-Delgado, L. & Espinosa, C.I. 2020. Knowledge gaps or change of distribution ranges? Explaining new records of birds in the Ecuadorian Tumbesian region of endemism. Diversity 12: 66. 10.3390/d12020066

Osorio-Olvera, L., Lira-Noriega, A., Soberón, J., Peterson, A.T., Falconi, M., Contreras-Díaz, R.G. & Barve, N. 2020. ntbox: an R package with graphical user interface for modelling and evaluating multidimensional ecological niches. Methods Ecol. Evol. 11: 1199–1206. 10.1111/2041-210X.13452

Paynter, R.J. 1997. Ornithological gazetteer of Colombia. 2nd edn. Museum of Comparative Zoology, Harvard University, Cambridge, MA.

Renjifo, L.M., Gómez, M.F., Velásquez-Tibatá, J., Amaya-Villarreal, A.M., Kattan, G.H., Amaya-Espinel, J.D. & Burbano-Girón, J. 2014. Libro rojo de aves de Colombia. Volumen I: bosques húmedos de los Andes y la costa Pacífica. Editorial Pontificia Universidad Javeriana & Instituto Alexander von Humboldt, Bogotá.

Ribas, C.C., Moyle, R.G., Miyaki, C.Y. & Cracraft, J. 2007. The assembly of montane biotas: linking Andean tectonics and climatic oscillations to independent regimes of diversification in Pionus parrots. Proc. R. Soc. B 274: 2399–2408. 10.1098/rspb.2007.0613

Rodríguez-Mahecha, J.V. & Hernández-Camacho, J.H. 2002. Loros de Colombia. Conservation International, Bogotá.

Salaman, P.G.W., Cuadros, T., Jaramillo, J.G. & Weber, W.H. 2001. Lista de chequeo de las aves de Colombia. Sociedad Antioqueña de Ornitología, Medellín.

Schindel, D.E. & Cook, J.A. 2018. The next generation of natural history collections. PLoS Biol. 16: e2006125. 10.1371/journal.pbio.2006125

Sullivan, B.L., Wood, C.L., Iliff, M.J., Bonney, R.E., Fink, D. & Kelling, S. 2009. eBird: a citizen-based bird observation network in the biological sciences. Biol. Conserv. 142: 2282–2292. 10.1016/j.biocon.2009.05.006

Watanabe, M.E. 2019. The evolution of natural history collections: new research tools move specimens and data to centre stage. BioScience 69: 163–169. 10.1093/biosci/biy163

